# OpenCRAVAT, an open source collaborative platform for the annotation of human genetic variation

**DOI:** 10.1101/794297

**Authors:** Kymberleigh A Pagel, Rick Kim, Kyle Moad, Ben Busby, Lily Zheng, Matthew Hynes-Grace, Collin Tokheim, Michael Ryan, Rachel Karchin

## Abstract

**PURPOSE:** The modern researcher is confronted with hundreds of published methods to interpret genetic variants. There are databases of genes and variants, phenotype-genotype relationships, algorithms that score and rank genes, and *in silic*o variant effect prediction tools. Because variant prioritization is a multi-factorial problem, a welcome development in the field has been the emergence of decision support frameworks, which make it easier to integrate multiple resources in an interactive environment. Current decision support frameworks are typically limited by closed proprietary architectures, access to a restricted set of tools, lack of customizability, web dependencies that expose protected data, or limited scalability.

**METHODS:** We present OpenCRAVAT, a new open source, scalable decision support system for variant and gene prioritization. We have designed the resource catalog to be open and modular to maximize community and developer involvement, and as a result the catalog is being actively developed and growing every month. Resources made available via the store are well-suited for analysis of cancer, as well as Mendelian and complex diseases.

**RESULTS:** OpenCRAVAT offers both command line utility and dynamic GUI, allowing users to install with a single command, easily download tools from an extensive resource catalog, create customized pipelines, and explore results in a richly detailed viewing environment. We present several case studies to illustrate the design of custom workflows to prioritize genes and variants.

**CONCLUSION:** OpenCRAVAT is distinguished from similar tools by its capabilities to access and integrate an unprecedented amount of diverse data resources and computational prediction methods, which span germline, somatic, common, rare, coding and non-coding variants. OpenCRAVAT is freely available at https://opencravat.org

## Introduction

Next-generation sequencing technologies have greatly reduced the cost of genome sequencing, increasing the availability of genomic data and the need for methods to evaluate genomic variants. The majority of variants have unclassified phenotypic consequences and their systematic exploration is complicated by data resources that are not easily obtainable or combinable. There is a need for more effective, user-friendly genome analysis tools that include interdisciplinary annotations and resources, to suit the needs of both novices and bioinformatics experts. Rapid identification of somatic variants relevant to the progression and treatment of cancer are of particular importance to facilitate timely precision patient care. Maintenance of patient privacy and data security place additional constraints on variant annotation and analysis, requiring systems that do not expose protected data.

Highly informative variant and gene characteristics are distributed across thousands of published works, spanning resources from the medical, biological, and bioinformatics domains, including experimental assays, computational variant effect prediction, evolutionary context, population databases, and established pharmacological relevance. This abundance of variant and gene annotations challenges researchers to broadly discover and deploy the best resources, as well as incorporate them within custom annotation pipelines. Furthermore, prediction algorithm software often requires nontrivial computational expertise to install, configure, and run. Recently genome-wide precomputation of predictor outputs for every possible input variant has been undertaken, to make computational tools more accessible. Databases that host precomputes, such as dbNSFP and WGSA ^1,2^ have been instrumental in exposing users to new tools. However, the datasets available from these resources were designed for machine rather than human access and require substantial programming investment before a user can incorporate them into an annotation pipeline.

Decision support framework (DSF) software tools have been created to integrate multiple annotation resources. Well-designed DSFs require substantial software development, and therefore the majority of DSFs are not freely available. The remaining minority of DSFs are either web-based portals that expose private data or downloadable tools with complicated installation and configuration requirements^3 4 5^. One such web-based DSF is CRAVAT (https://cravat.us), which prioritizes somatic mutations^6^. In this work, we present OpenCRAVAT, an extension of CRAVAT with improved data security, a much larger collection of annotations, and the capability to generate dynamic and customizable pipelines.

The Open Custom Ranked Analysis of Variants Toolkit (OpenCRAVAT) is a freely available open source framework for the annotation and visualization of human genetic variation and genomic elements. The framework can rapidly generate publication quality visualizations of gene networks, the distribution of variants per protein, and supports BAM file visualization with an embedded version of the Integrative Genomics Viewer (IGV) ^*7*^. Designed to comprehensively annotate both well-characterized and novel somatic and germline variation, the framework can be flexibly adapted to suit a wide spectrum of human variation research projects. In this work, we describe the underlying architecture and present several use cases.

## Methods

### Framework architecture

OpenCRAVAT is written in Python and all code is stored on a public repository. It is open source and free of charge to users, with both command line and graphical user interface functionality. OpenCRAVAT can be installed via user-friendly wizard or through pip. The framework is built around two main components: a base module and a store where users can download additional modules. Modules include input format Converters, gene Mappers, Annotators, output format Reporters, and graphical Widgets. The base module includes converters that support Variant Call Format (VCF), tab-delimited (TSV) and comma-delimited (CSV) text files, a mapper that projects genome positions to transcript, protein sequence and protein structure coordinates, a set of basic Widgets and Reporters that generate results in sqlite3, Excel, TSV, and CSV formats. OpenCRAVAT supports GRCh38, GRCh37 and GRCh36 human genome reference assemblies, and variants are mapped to all GENCODE isoforms^8^. The store offers a large selection of modules, including additional Converters (PLINK, Ancestry, 23andMe, dbSNP identifiers), Annotators for somatic, de novo and germline variation (coding and noncoding), associated Widgets and Reporters (VCF, pipeline-friendly TSV and CSV).

The store is available through both GUI and command-line interface. Within the GUI, available modules are displayed in a format similar to an app store, where each tool is represented by a tile containing documentation, update status, and one-click installation. After installation, OpenCRAVAT downloads each resource locally, which enables secure analysis of private data. The open store is built for continuous community-driven development, so that newly developed tools and resources can be uploaded and made available to a wide audience. Addition of new resources to the store requires data descriptions, appropriately formatted annotation data, and a small script to allow incorporation of the data by OpenCRAVAT. Module developers can select whether to openly publish their data or restrict access with the option to share the Module directly with appropriate collaborators.

### Using OpenCRAVAT

Configurable workflows within OpenCRAVAT can be created and executed in either the command-line or graphical user interface. OpenCRAVAT generates annotations for input files of human genetic variants. VCF, annotated VCF, basic tabular file format, dbSNP identifiers, 23andMe and Ancestry.com files are supported. To accommodate family and cohort studies, multiple VCF files can be selected and merged within a single annotation run, in addition to support for multi-sample VCF files. For each annotation run, the user has the option to include all installed annotators or a subset, allowing for the creation of custom annotation pipelines (Figure 1). Upon completion of a run, the interactive results viewer can be used for exploratory data analysis and filtering.

**Figure 1.**
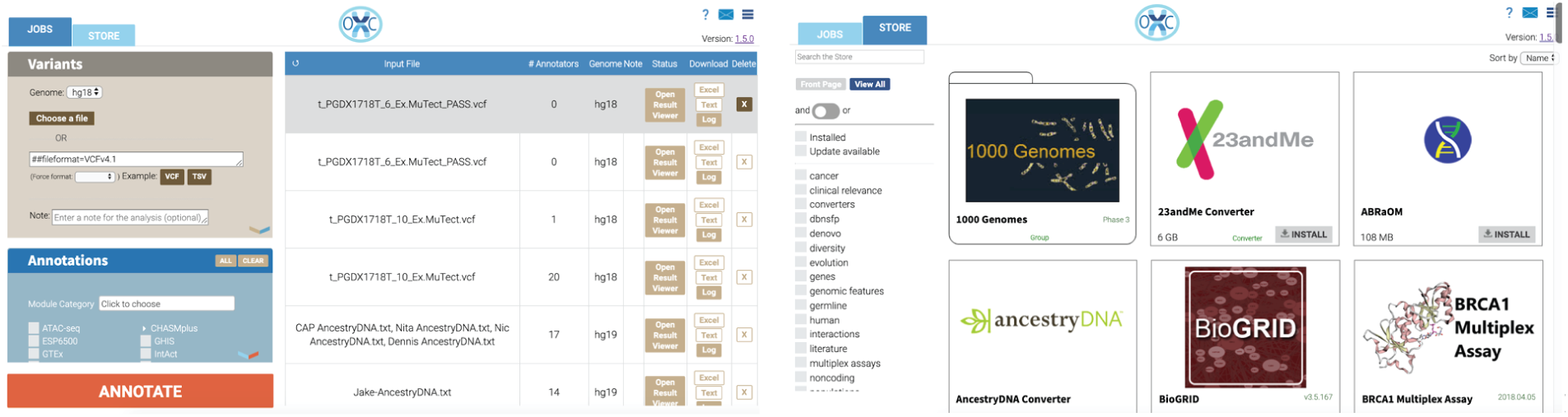
Screenshot of the OpenCRAVAT GUI submission and store pages.

Accessible via both command line and graphical user interface, the viewer comprises four tabs: Summary, Variant, Gene, and Filter (Figure 2). The Summary tab displays graphical representation of the submitted variant characteristics, as well as submission details including the selected annotations and data source versions. The Variant and Gene tabs are divided into an interactive table and widget pane. The interactive table displays each variant or gene on a particular row along with the corresponding user-selected annotations. The widget pane includes several interactive elements which graphically display further information and visualizations of the annotators including the Integrative Genomics Viewer (IGV) with BAM file support and a Protein Diagram to visualize protein-level variation. Within the viewer, table columns and widgets can be resized or hidden, and layout preferences can be saved, shared, and applied to other annotation runs. The Filter tab allows to users generate and save filters, which identify variants in selected samples or genes, population allele frequency ranges, genomic locations, by sequence ontology, or custom annotator-specific thresholds. For more complex filtering, the graphical Query Builder can build advanced SQL queries.

**Figure 2.**
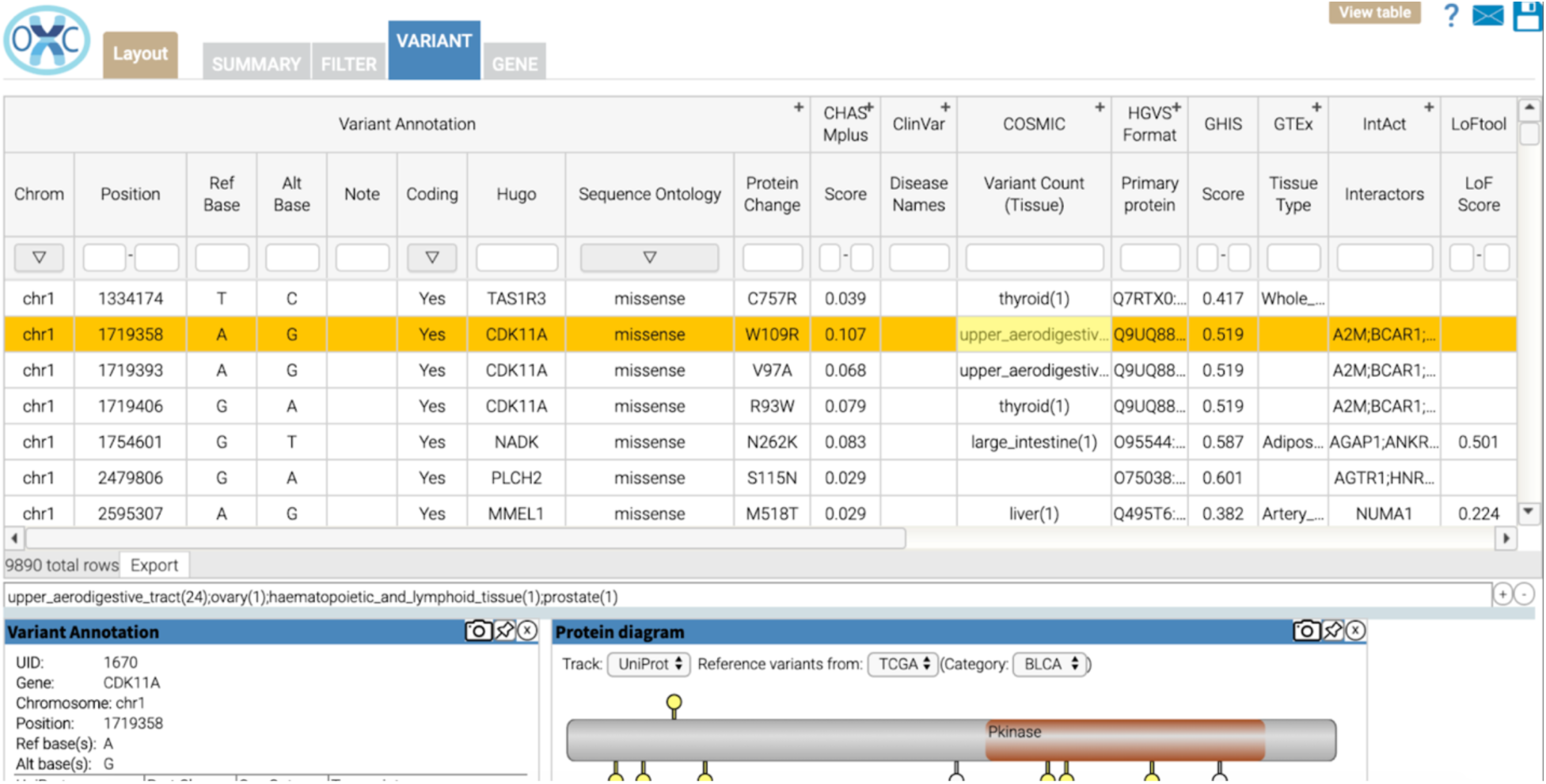
Screenshot of the OpenCRAVAT GUI variant analysis table.

OpenCRAVAT can be installed locally on a user’s computer or on a server, allowing multiple users to submit annotation runs on the same system, with administrator monitoring and maintenance. The server implementation adds user authentication, user-specific storage, user access to history and shared access to analysis and visualization results. Server installation can be performed on both a shared local system or in a cloud environment, where results storage can be controlled and protected data is secure. The entire catalog of resources can be stored in one place and shared among many users, in addition to analysis results.

## Results

In the following case studies, we illustrate the capacity of OpenCRAVAT to evaluate phenotypically relevant genetic variations within inputs of differing size and composition.

### Case Study 1: Variant prioritization in multiple lesion cancer samples

Among the somatic variants present in a tumor, a small number of mutations are believed to “drive” tumor growth, and may be useful for diagnosis, prognosis, patient stratification, clinical trial eligibility and selection of appropriate therapies. Of particular interest are clonal driver mutations which occurred in the initiating tumor cell and are present in all tumor cells. Identification of these originating mutations can be enhanced by evaluating mutations from multiple tumor biopsies, including precursor lesions, primary cancers and metastases from a single patient.

In this case study, we investigated early candidate driver mutations in a high-grade serous ovarian cancer patient (CGOV62), using VCF and BAM files from a published genomic study of high-grade serous ovarian cancers, including fallopian tube precursor lesions, fallopian tube and ovarian tumors and omental, rectal and appendiceal metastases, and a normal fallopian tube epithelium control sample ^9^. BAM files from whole-exome sequencing were downloaded from the European Bioinformatics Institute (EGAS00001002589), VCF files were generated with MuTect v.1.1.7 using default parameters^10^.

The analysis was carried out using the Query Builder (Figure 3A):

**Figure 3:**
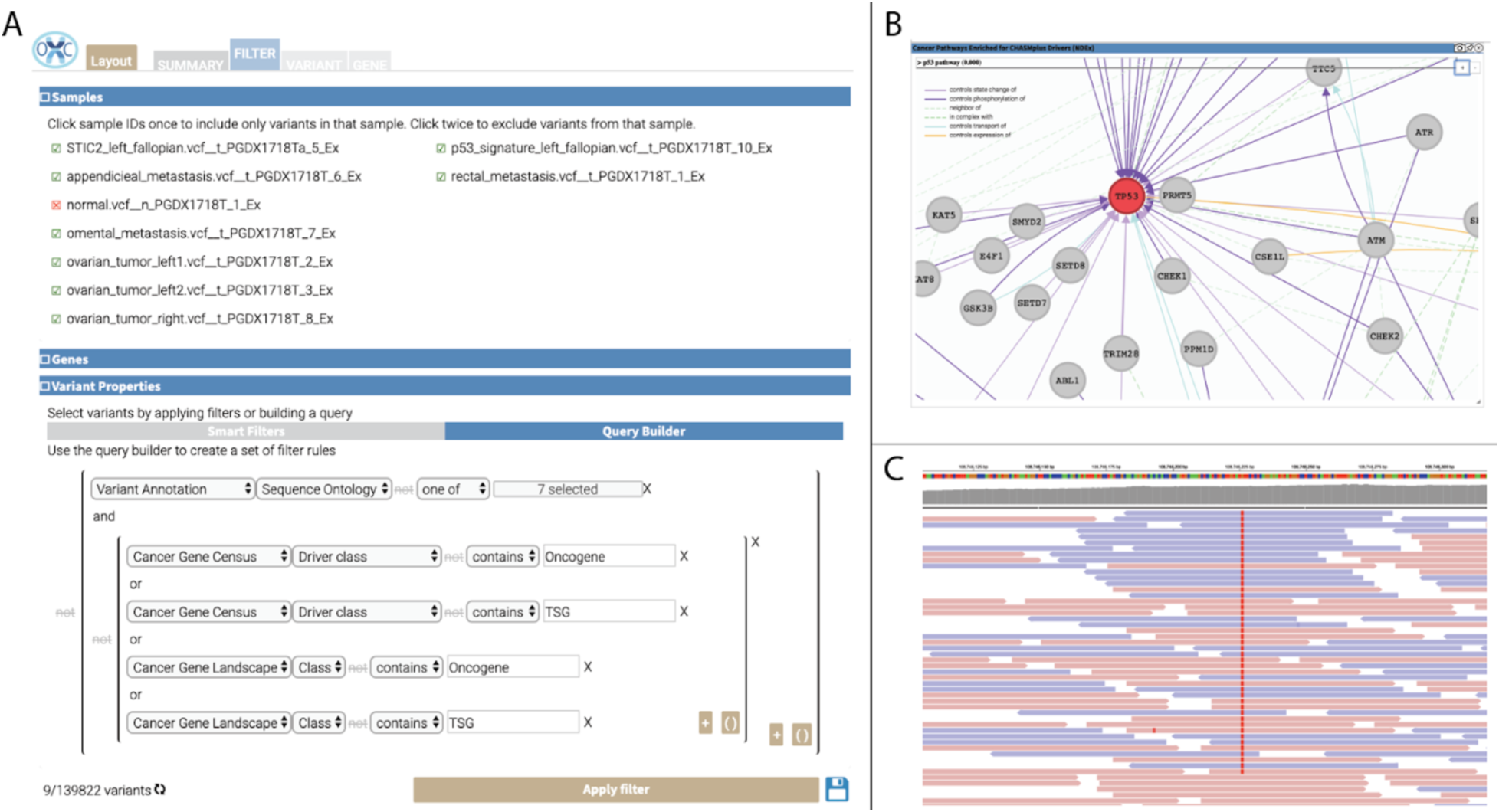
Components of the OpenCRAVAT GUI used in Case Study 1. (A) The Query Builder filters applied to identify potential cancer driver mutations. (B) NDEx network enriched for mutations within these samples. (C) Screenshot of IGV reads for a tumor sample.

1. Installed cancer-related annotation modules (*Cancer Gene Census^11^*, *Cancer Gene Landscapes^12^*), computational predictors (*CHASMplus OV^13^*, *MutPred^14^*) and a visualization module (*IGV*).
2. Within the interactive interface, selected the genome version used in the study (hg18), uploaded VCF files for each biopsied lesion, selected the annotators listed above, and clicked the Annotate button.
3. On the Filter tab, filtered by sample to exclude any germline variants that were present in the normal fallopian tube epithelium sample.
4. To focus on loss-of-function mutations in tumor suppressor genes (TSG) and missense mutations in oncogenes (OG), applied a Sequence Ontology filter to select either (missense, splice site, frameshift and non-frameshift indels, and stop gain) for TSG or (missense and non-frameshift indels) for OG.
5. Retained mutations within known OG and TSG as provided by either the Cancer Gene Landscapes or Cancer Gene Census.

Nine mutations were retained after applying these filters, of which two are likely clonal mutations: *RANBP2*:p.M933I and *TP53*:p.T126N. These mutations were observed in seven of the eight lesions. In the original study, the *TP53* mutation was found in an eighth lesion by deep targeted sequencing. The *TP53* mutation is a known driver, with CHASMplus OV p-value less than 0.01, and is predicted by MutPred to result in loss of sheet structure (p-value=0.0457). The NDEx Widget was used to explore interaction partners of the mutated proteins, and the NDEx enrichment tool identified 13 *TP53*-associated networks from the NCI Pathway Interaction Database^15^ (Figure 3B). For each truncal mutation, the normal and tumor BAM files were loaded into *IGV* for viewing and manual validation (Figure 3C). Manual inspection verified that the mutation was truly somatic, not present in normal tissue (data not shown), and that there was no apparent strand bias.

### Case study 2: Identifying driver missense mutations among metastases

We analyzed exome and genome sequencing data for 76 untreated metastases from 20 patients with breast, colorectal, endometrial, gastric, lung, melanoma, pancreatic, and prostate cancers from a recent study on the heterogeneity of functional driver mutations in cancer metastases^16^.

1. Installed the *CHASMplus* Annotator to score mutations as likely cancer drivers and *tsvreporter* to generate simple tab-delimited output.
2. Assembled a tab-delimited file of 15,765 somatic mutations identified in the Reiter et al. study: *reiter_et_al_2018.txt* (Data S1).
3. Used the command line interface to generate a CHASMplus score for each mutation:

~~~
cravat reiter_et_al_2018.txt -n Reiter_2018 -t tsv -l hg19 -
-cleanup -d output
~~~
4. Ran a python script *fdr.py* that took in the output file and created a qvalue for each mutation, which is a correction of the CHASMplus p-value for multiple hypothesis testing using the False Discovery Rate of 0.05 (Data S2).

In total, 56 mutations were predicted as drivers, with a significant qvalue (q<0.01). These included well-known oncogenic alleles (*KRAS*:p.G12D, *SMAD4*:p.D351G) and *PTEN*:p.R173H. A literature search confirmed that *PTEN*:p.R173H was identified as functionally damaging in saturation mutagenesis assays^17^.

### Case study 3: Clinically actionable variants an individual genome

To identify variants with potential for clinical relevance, we applied filters to the whole genome sequence variation of a phenotypically normal individual obtained from the Personal Genome Project (Profile hu3BDC4B, https://www.personalgenomes.org) ^18^. First, we used guidelines from the American College of Medical Genetics (ACMG)^19^ to classify variants with “Very strong evidence of pathogenicity”. These guidelines define Very strong evidence of pathogenicity, as “Null variant (nonsense, frameshift, canonical +/−1 or 2 splice sites, initiation codon, single or multi-exon deletion) in a gene where loss of function (LOF) is a known mechanism of disease”. To replicate the guideline, we identified putative disease-relevant haploinsufficient genes, using LoFtool, a gene intolerance ranking system^20^ and a curated list of gene-disease associations from ClinGen^21^.

1. Installed *LoFtool*, *gnomAD* and *ClinGen Gene*.
2. Within the interactive interface, selected the genome version used in the study (hg19), uploaded the VCF and clicked the Annotate button.
3. On the filter tab, retained variants categorized by the base *Sequence Ontology* annotator as “stop gained”, “frameshift deletion”, “frameshift insertion”, or “splice site”.
4. Excluded variants within genes with LoF Score<0.9
5. Excluded 19 variants with gnomAD Global allele frequency >0.05 which is considered “Stand alone” evidence of Benign impact by the ACMG Guidelines.
6. Retained variants within genes with known disease association.

Out of 3,926,112 variants in this genome, two variants met these criteria and impact two genes (*PALB2* and *PCDH15)*. This example illustrates how OpenCRAVAT can be used to reproduce a variant filtering workflow which previously has required development of dedicated software.

### Case Study 4: Identification of variants that segregate with disease within a quartet

To identify genetic loci segregating with disease using variation from four members of a nuclear family, we obtained genotyping information of the family from Ancestry.com. Within the extended family pedigree (Figure 4A), two impacted family members, the proband and maternal uncle, exhibited both keratoconus and dyslexia/dysgraphia.

**Figure 4.**
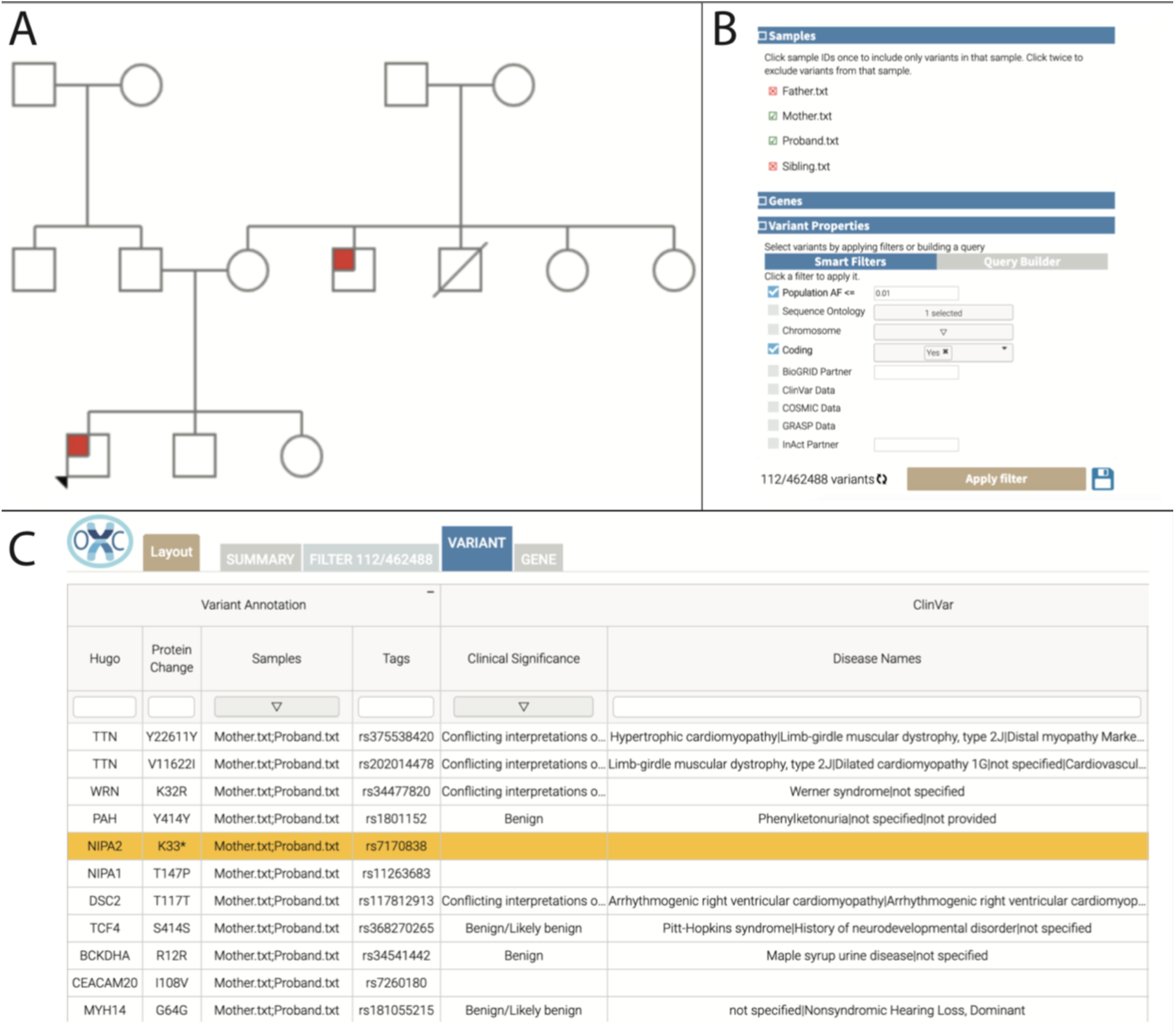
(A) Family pedigree. The proband is shown at the lower left in red, and an affected maternal uncle is designated by the remaining red square. (B) Filters used to identify potentially relevant variants. (C) The 11 low frequency variants present in both Proband and Mother.

1. Installed modules to identify disease-causing variants (*ClinVar^22^*), allele frequency annotations (*gnomAD*, *1000 Genomes Projec*t^23^), *LoFtool*, and the Ancestry.com input converter (*AncestryConverter*).
2. Within the interactive interface, selected the genome version used in the study (hg19), uploaded files for Unaffected carrier mother, father, unaffected male sibling, and the proband (Personal Genome Project Profiles hu58B346, huB08B19, hu6A2DBA, huB1557E) and clicked the Annotate button.
3. Applied sample-level filters to retain only variants that agree with the inheritance pattern observed in the family: Variants within the father and unaffected sibling were removed, assuming that neither carried the causative variant due to lack of phenotype (Figure 4B).
4. Assuming that these variants are rare, removed variants with Global Allele Frequency > 0.01 in gnomAD.
5. Assessed each remaining variants for previously established phenotypic relevance in ClinVar.

Eleven low-frequency variants were identified in both Proband and Mother, of which four are “Benign” in ClinVar, and four have “Conflicting Interpretations” (Figure 4C). The remaining three variants were *CEACAM20*:p.I108V, *NIPA1*:p.T147P, and *NIPA2*:p.K33*.

NCBI Gene summary widget showed that *NIPA1* and *NIPA2* are known/putative magnesium transporters. *NIPA2* is predicted to be intolerant for loss-of-function by LoFtool score of 0.054, supporting a hypothesis for the phenotypic relevance of *NIPA2*:p.K33*. Manual literature search revealed a microdeletion syndrome that includes the loss of *NIPA1* and *NIPA2* and carries a high risk of reading and writing difficulties^24^, further supporting the possibility that these genes play a role in the observed phenotype.

### Case Study 5: Identification of dbSNP entries in linkage disequilibrium with pathogenic variants

In collaboration with the University of Colorado, we are using OpenCRAVAT on Amazon Web Services (AWS) to annotate every available rsID in dbSNP build 153 and identify all SNPs in close proximity to pathogenic variants listed in ClinVar. Here, we describe performance of the workflow on chromosome 20. The analysis consisted of the following steps:

1. Created an Amazon Machine Image (AMI) of OpenCRAVAT.
2. Installed all available modules to ensure the most comprehensive possible annotations. In particular, disease-causing variants (*ClinVar*), dbSNP input converter (*dbSNPConverter*) and linkage-disequilibrium (*LDAnnotate*).
3. To parallelize the analysis, a Cloud Formation (CF) workflow was used to process dbSNP rsIDs by chromosome across multiple instances of the OpenCRAVAT AMI. Input and config files from S3 were passed to a CF template which automated the processing and results delivery.

The 14,663,863 SNPs on chromosome 20 were processed in 14.5 hours. For each pathogenic ClinVar variant with a dbSNP identifier, we identified all dbSNP entries that are in linkage disequilibrium (D’ > 0.5). The OpenCRAVAT AMI, CF template, input, config and results files are available at https://opencravat.org/resources.html.

## Discussion

OpenCRAVAT is a flexible and dynamic system to annotate and evaluate the characteristics of genetic variation. It has been designed to enable rapid characterization of variants, including functional impact, pharmacological annotations, and both known and predicted relevance of genetic variants to disease, including cancer. The open store contains dozens of resources relevant to variant interpretation, with new additions weekly. Selection of specialized converters, annotators and filtering criteria enable researchers to carry out complex analyses and integrate information from a wider array of resources than previously possible.

We have described a framework that includes both an advanced GUI for biologists and a command line interface that supports advanced use cases, including development of custom bioinformatics pipelines. Both GUI and command line can be leveraged in the cloud to handle processing of genomes from large patient populations. Finally, as OpenCRAVAT is designed to be community-driven we have incorporated 68 tools from 37 universities/institutes in the past year, and we are actively recruiting tool and resource developers.

## Availability

https://github.com/KarchinLab/open-cravat/

